# Circulatory neutrophils exhibit enhanced neutrophil extracellular trap formation in early puerperium: NETs at the nexus of thrombosis and immunity?

**DOI:** 10.1101/2021.10.18.459827

**Authors:** Stavros Giaglis, Chanchal Sur Chowdhury, Shane Vontelin van Breda, Maria Stoikou, Guenther Schaefer, Andreas Buser, Ulrich A. Walker, Olav Lapaire, Irene Hoesli, Paul Hasler, Sinuhe Hahn

## Abstract

Pregnancy is associated with elevated maternal levels of cell-free DNA of neutrophil extracellular trap (NET) origin, as circulatory neutrophils exhibit increased spontaneous NET formation, mainly driven by G-CSF and finely modulated by sex hormones. The postpartum period, on the other hand, involves physiological alterations consistent with the need for protection against infections and fatal haemorrhage. Our findings indicate that all relevant serum markers of neutrophil degranulation and NET release are substantially augmented postpartum. Neutrophil pro-NETotic activity *in vitro* is also upregulated particularly in post-delivery neutrophils. Moreover, maternal puerperal neutrophils exhibit a strong pro-NETotic phenotype, associated with increased levels of all key players in the generation of NETs – citH3, MPO, NE and ROS, compared to non-pregnant and pregnant controls. Intriguingly, post-delivery NET formation is independent of G-CSF in contrast to late gestation and complemented by the presence of TF on the NETs, alterations in the platelet activity status, and activation of the coagulation cascade, triggered by circulating microparticles. Taken together, our results reveal the highly pro-NETotic and potentially procoagulant nature of postpartum neutrophils, bridging an overt immune activation with possible harmful thrombotic incidence.

## Introduction

The postpartum period involves physiological alterations in the response of cellular blood components, consistent with the need for extensive protection against potential infections and haemorrhage required in this phase (1). Recent evidence implies that the activation of the innate immune system during and shortly after childbirth is functionally essential for endogenous repair processes (2, 3) and that an excessive immune response might be detrimental for the well-being of both the mother and child, eventually with even fatal consequences (4), as postpartum haemorrhage, puerperal infections and venous thromboembolism (VTE) are still related to high maternal mortality and morbidity (1, 5, 6). Therefore, fast deployment of antimicrobial defence mechanisms, prompt activation of the coagulation cascade and timely initiation of an effective wound healing sequence are of great relevance. is required.

Neutrophilic granulocytes are involved in the first line of antimicrobial defence (7, 8). In addition to playing an important role in direct bacterial killing by phagocytosis and degranulation, neutrophil activation leads to the release of nuclear and granular contents to form web-like structures of DNA, coined neutrophil extracellular traps (NETs) (9, 10). The extracellular DNA is decorated with nucleosomal material – histones – and a series of potent granular enzymes including neutrophil elastase (NE), cathepsin G, and myeloperoxidase (MPO) (11). These structures have also been recently identified in patients with autoimmune disorders (12, 13), inflammation (14), cardiovascular and pulmonary diseases (15, 16) and thrombosis (17, 18). Blood-derived neutrophils and release of NETs have been identified in animal models of recurrent fetal loss and in placental tissue of patients with preeclampsia (19, 20). In pregnancy the regulation of neutrophil function and NET formation is driven predominantly by sex hormones and circulatory granulocyte colony-stimulating factor (G-CSF), and also associated with an increased risk for thrombosis (21, 22). The function of neutrophils during gestation and especially the early postpartum period is poorly studied. Furthermore, whether or not NETs contribute to favourable or poor outcomes after term and recovery remains yet unclear.

We and others have previously reported that the levels of maternal cell-free DNA are elevated in the serum of pregnant women, especially in cases affected by preeclampsia (23, 24). It was subsequently verified that neutrophil NETs contribute to the circulating pool of this material (14, 20, 21). In the present study, we detect elevated levels of circulating DNA in early postpartum serum and demonstrate the enhanced tendency of neutrophils to form NETs *in vitro* which culminates 48-hours after delivery. We also reveal that this process is independent of G-CSF, possibly the most potent regulator of neutrophil activity in general, and that post-delivery NETs are decorated with TF. Finally, we describe the generation of NETs as a response to plasma microparticles by ways of a secondary hit, complementing thereby the antimicrobial and procoagulant propensities of postpartum neutrophils.

## Materials and methods

### Human Subjects

Pregnant women were recruited at the time of their routine examination at the end of the first (median gestational age: 12 weeks and 11 days - n=5; median age: 34 years) and second trimesters (median gestational age: 24 weeks and 7 days - n=13; median age: 36 years) and at the time of elective Caesarean section at the end of the third trimester (median gestational age at delivery: 38 weeks and 12 days - n=11; median age: 39 years); part of the individuals from the third trimester group (n=9) donated blood samples 48 hours after delivery. Healthy volunteers, matched for gender and age (n=20; median age: 32 years), were recruited at the Blood Bank of the Swiss Red Cross, Basel. Inclusion criteria for healthy controls were fair general condition, age ≥ 25 and ≤ 45 years and for blood donors fulfilling national criteria for blood donation. All blood donor data are summarized in Supplementary Table S1. Exclusion criteria were current or previous systemic autoimmune disease, asthma, reconvalescence after major illness, surgery, current medication with corticosteroids, immunosuppressive agents and malignant neoplasia or chemotherapy within 5 years before recruitment for the study. Exclusion criteria included any major complication of pregnancy or coincident disease, such as pre-eclampsia, pre- or post-term labor (<37 weeks or >42 weeks), intra-uterine growth retardation and viral, bacterial or parasitic infections. Informed, written consent was obtained from all subjects in the study, which was approved by the Ethical Review Board of Basel/Basel-Land, Switzerland.

### Preparation of plasma and serum

Plasma and serum was collected and processed as described previously (25). Samples were studied immediately or stored at -80°C until analysis. Microdebris was pelleted by two consecutive centrifugation steps at 20,000xg for 15 minutes at 4°C. Microparticle-depleted plasma (supernatant) was verified by flow cytometry as previously described (26).

### Blood cell count and preparation of plasma and serum

Whole blood was collected into EDTA- and silicone-coated tubes (Sarstedt) and 25μl of blood was analyzed by a Hemavet 950FS (Drew Scientific) for complete blood cell counts. Plasma and serum was collected and processed as described previously (25). All collected samples were studied immediately or stored at -80°C until analysis.

### Human neutrophil isolation

PMNs were isolated by Dextran-Ficoll density centrifugation (12, 21). Briefly, EDTA-containing blood was diluted with PBS/EDTA 2mM (Gibco Life Technologies), and layered on a Ficoll Paque plus gradient (GE) in order to deplete the mononuclear cells fraction. After centrifugation, neutrophils were enriched by separation from the erythrocyte pellet through Dextran sedimentation (Sigma). The contaminating erythrocytes were lysed in ice-cold RBC lysis buffer (Biolegend). PMNs were rinsed twice and re-suspended in Hanks’ balanced salt solution (HBSS; without calcium/magnesium and phenol red) (Gibco Life Technologies) supplemented with 2% autologous plasma. Cell viability was assessed by trypan blue dye exclusion in a haemocytometer and was routinely 96– 98% with a purity of > 95% PMNs. PMNs were directly seeded in 24-well or 96-well plates and allowed to settle for 15 minutes at 37°C under 5% CO_2_ prior to further experimentation. For examining spontaneous NETosis *in vitro*, a time-course of two time-points after 15 minutes of settling was chosen: a baseline initial condition (0h) and a 3-hour culture period (3h).

### Cell free DNA isolation and quantification

Cell free DNA was extracted from 850μl serum using the QIAamp Circulating Nucleic Acid Kit (Qiagen) and was quantified by TaqMan Real-time PCR (StepOneTM Plus Real-Time PCR System, Applied Biosystems) specific for the glyceraldehyde-3-phosphate dehydrogenase (*GAPDH*) gene (25).

### Stimulation and neutralization studies

For *in vitro* incubation studies 2.5 × 10^4^ PMNs from control healthy individuals were treated with 3% serum or 8% plasma derived from control non-pregnant individuals, pregnant donors during the first, second and third trimester of gestation; similarly, 48 hours and one month post-delivery donor serum was used. All experiments were carried out in a 3 hours timecourse. To neutralize serum G-CSF, pooled plasma or depleted plasma from the study groups of interest were pretreated with anti-G-CSF Ab (0.2 μg/ml, Peprotech) for 30 minutes.

### Microparticle isolation and characterisation

Circulating microparticles were isolated, characterised and utilized as described previously (18).

### Fluorimetric quantification and fluorescence microscopy

NETs were quantified using SytoxGreen fluorimetry as described previously (19, 27, 28). Briefly, 2.5×10^4^ freshly isolated neutrophils were cultured in the presence of 0.2 μM SytoxGreen (Invitrogen Life Technologies) in a 96-well dark microtitre plate at 37°C under 5% CO2 and either left untreated or stimulated with the aforementioned stimuli or PMA as a positive control for the indicated 3-hour timecourse. Fluorescence (excitation 485 nm, emission 535 nm) was measured in a Biotek Synergy H1 Hybrid Reader (Biotek) and results reported as DNA fluorescence (MFI). Photomicrographs in brightfield and green fluorescence spectra were assessed with an Olympus IX50 inverted fluorescence microscope (Olympus) coupled to an Olympus XM10 monochromatic CCD camera (Olympus) and analyzed with the Olympus CellSens Dimension software (Olympus).

### Neutrophil elastase (NE), myeloperoxidase (MPO), cell-free histone/DNA complex, MPO/DNA complex, thrombin-antithrombin (TAT) complex and D-dimers analysis

Protein quantification analysis was performed by ELISA assays, as previously described (21). To detect NET-associated MPO/DNA complexes, a modified capture ELISA was utilized (13). The concentrations of thrombin-antithrombin (TAT) complexes and D-dimers were measured by sandwich ELISA, utilizing the human TAT Complexes ELISA Kit (Assaypro) and the Imuclone D-Dimer ELISA Kit (American Diagnostica) respectively.

### RNA Isolation and Quantitative Real-time PCR

Total RNA was isolated from 3×10^6^ neutrophils by using the RNeasy Mini Kit (Qiagen). TaqMan real-time quantitative RT-PCR was performed utilizing the Applied Biosystems StepOne Plus cycler (Applied Biosystems) and TaqMan Gene Expression Assay primer and probe sets (Applied Biosystems) for ELANE (HS00236952_m1). Data were normalized to the housekeeping gene B2M (HS99999907_m1), after a selection procedure from six different endogenous reference genes, as suggested in the MIQE guidelines (32). Relative values were calculated with 2-DDCt analysis (33).

### Oxidative burst analysis

NADPH oxidase mediated ROS production was measured by either using a 2′,7′-dichloro dihydrofluorescein diacetate (DCFH-DA) plate assay (29) or a luminol-based chemiluminescence microtitre plate assay, as previously described (30).

### Fluorescence microscopy

5 × 10^4^ isolated PMNs were seeded and incubated for 10min with 5µM Sytox Green dye (Invitrogen Life Technologies) for assessment of NETs with an Axiovert fluorescence microscope coupled to a Zeiss AxioCam colour CCD camera (Carl Zeiss) (19).

### Immunohistochemical microscopy and morphometric analysis

NETs were quantified after IHC staining of 2.5 × 10^4^ PMNs per well in a 96-well plate with mouse anti-human MPO antibody (Abcam) and rabbit anti-human citH3 antibody (Abcam), or the respective isotype controls, followed by incubation with goat anti-mouse IgG AF555 and goat anti-rabbit IgG AF488 (Invitrogen Life Technologies). DNA was counterstained with 4′,6-diamidino-2-phenylindole (DAPI, Sigma-Aldrich). NETs were visualized by using an Olympus IX81 motorized epifluorescence microscope (Olympus America Inc., Center Valley, PA, USA) in conjunction with an Olympus XM10 monochromatic CCD camera (Olympus) and analyzed with the Olympus CellSens Dimension software (Olympus). A minimum of 20 fields at 10× magnification (at least 500 to 1,000 PMNs) per case was evaluated for MPO/citH3 and DNA co-staining through ImageJ analysis software (National Institutes of Health Image Processing, Bethesda, MD, USA); nuclear phenotypes and NETs were determined, counted, and expressed as percentage of the total area of cells in the fields (31).

### Phagocytosis activity assessment

Neutrophil phagocytic activity was examined by measuring the amount of uptake of latex beads coated with FITC-labeled rabbit IgG into cells using a phagocytosis assay kit (Cayman Chemical) as previously described (21).

### Protein isolation and western blot analysis for PAD4, citH3, NE and MPO

Western blot analysis was performed as previously described (28). Briefly, total protein was isolated by NucleoSpin TriPrep kit (Macherey-Nagel) from 5×10^6^ PMNs. All protein concentrations were determined with the MN Protein Quantification Assay (Macherey-Nagel). Western blotting was performed utilizing AnykD Mini-PROTEAN TGX Gels (Biorad) and nylon/nitrocellulose membranes (Biorad). Equal loading was verified using beta-actin. Western blots of citrullinated H3 (citH3) protein were prepared as described previously (32). Gel documentation, densitometric analysis and protein quantification of the western blots was performed using the ChemiDoc XRS+ maging system (Biorad) with the ImageLab 4.1 image analysis software (Biorad).

### Microparticle isolation and characterisation

Circulating microparticles were isolated, characterised and utilized as described previously (18).

### Statistical analysis

All data are presented as mean ± SE. Descriptive statistics for continuous parameters consisted of median and range, and categorical variables were expressed as percentages. Comparisons between patients and healthy controls were carried out by the Mann-Whitney U test with a Welch post-test correction. Statistical significance in multiple comparisons was by one-way analysis of variance (ANOVA) with a Dunn’s post-test correction. P values under < 0.05 were considered as statistically significant. Data were processed in GraphPad Prism version 9.0 for MacOSX (GraphPad Software Inc., www.graphpad.com).

## Results

### Serum markers of neutrophil degranulation and NETosis are augmented in early puerperium

In this study cohort’s serum samples, all markers of neutrophil degranulation and NETosis were found to be augmented post-delivery. In particular, real-time PCR and dsDNA-binding fluorescent dye quantification revealed elevated cell-free DNA levels in these sera (Figure 1a), and the nucleosomal nature of the detected DNA was additionally verified by nucleosome specific ELISA (Figure 1b). This was shown to be complexed with myeloperoxidase (MPO), a potent lysosomal protease stored in the azurophilic granules of the neutrophil (Figure 1b), indicating that the cell-free DNA is of NETotic origin (13). In contrast to the serum, none of the plasma levels of these NET components attained statistical significance, signifying the strong reactivity of postpartum neutrophils during the blood clotting process. Additionally, serum MPO and neutrophil elastase (NE) levels were also elevated postpartum (Figure 1c), the latter, a potent serine protease secreted by neutrophils during inflammation. Interestingly, pronounced neutrophilia and concomitant decrease in monocyte numbers were more pronounced post-delivery compared to the non-pregnant controls, possibly indicating a distinct biological mechanism driving the release of NET (Figure 1d).

**Figure 1.**
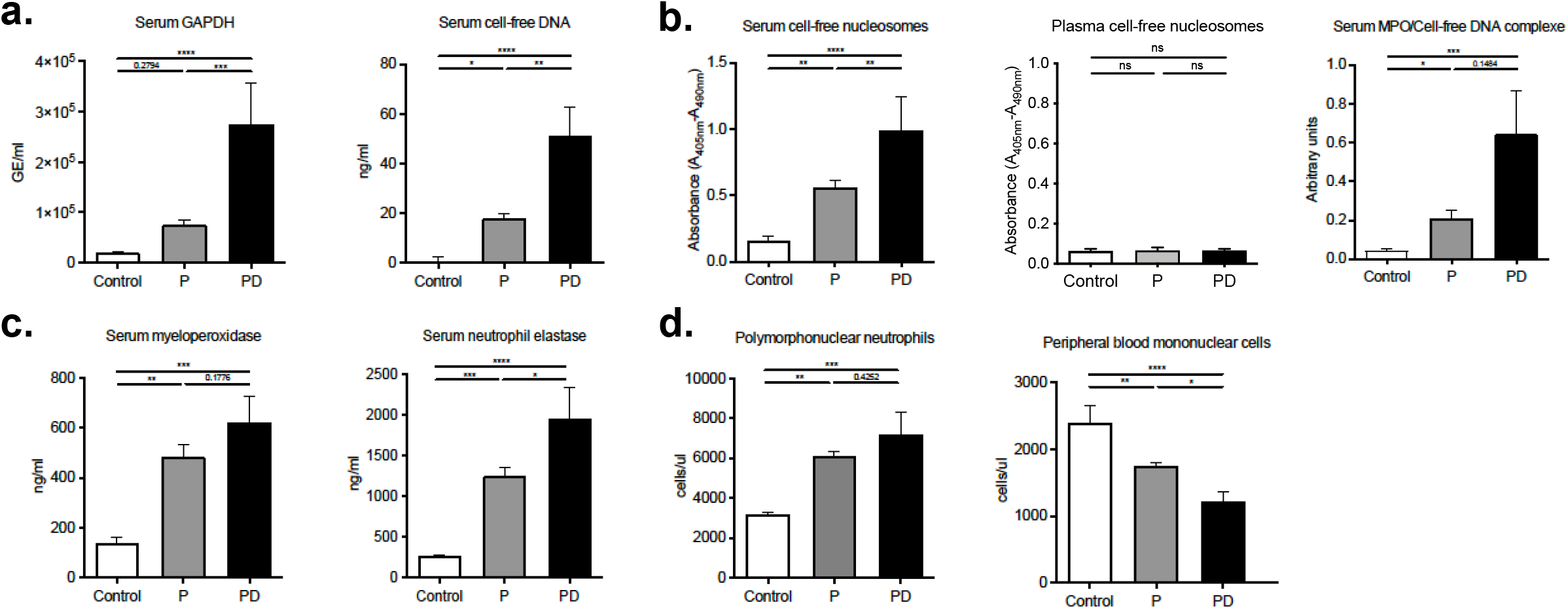
Serum markers of neutrophil degranulation and NETosis are substantially increased 48 hours post-delivery. (**a**.) Cell-free DNA levels in serum from donors during pregnancy (n=12), 48 hours postpartum (n=9) and healthy matched blood donors (n=11), determined by GAPDH real-time PCR (left panel) and dsDNA-binding fluorescent dye quantification (right panel). (**b**.) Cell-free nucleosome levels in serum from healthy donor controls, donors during gestation and postpartum, determined by ELISA and NET-associated MPO/DNA complexes quantified utilizing a modified capture ELISA. In contrast to the serum levels, none of the plasma levels of these NET components attained statistical significance. (**c**.) MPO and NE concentrations in serum from healthy donors, donors during pregnancy and 48 hours postpartum as determined by sandwich ELISA. (**d**.) Neutrophil and peripheral blood mononuclear differential cell counts in healthy non-pregnant blood donors, donors during pregnancy and 48 hours postpartum. P: pregnancy; PD: post-delivery. Data are presented as mean ± SEM. ^*^P < 0.05, ^**^P < 0.01, ^***^P < 0.001, ^****^P < 0.0001.

This finding prompted us to test whether postpartum neutrophils indeed generate spontaneously cell-free NETs. For this purpose, we performed an *in vitro* 3-hour culture of isolated neutrophils where enhanced NET formation by pregnancy and postpartum-derived neutrophils was observed (14, 33) compared to the control neutrophils (Figure 2a). In this preliminary experiment, NETs were detected by simple fluorescence detection of SytoxGreen staining of the extracellular DNA structures (19, 33). In parallel, the levels of intra-and extracellular ROS (Figure 2b and 2c), as well as the phagocytic activity of the postpartum neutrophils (Figure 2d) was markedly elevated compared to both non-pregnant control and pregnancy neutrophils.

**Figure 2.**
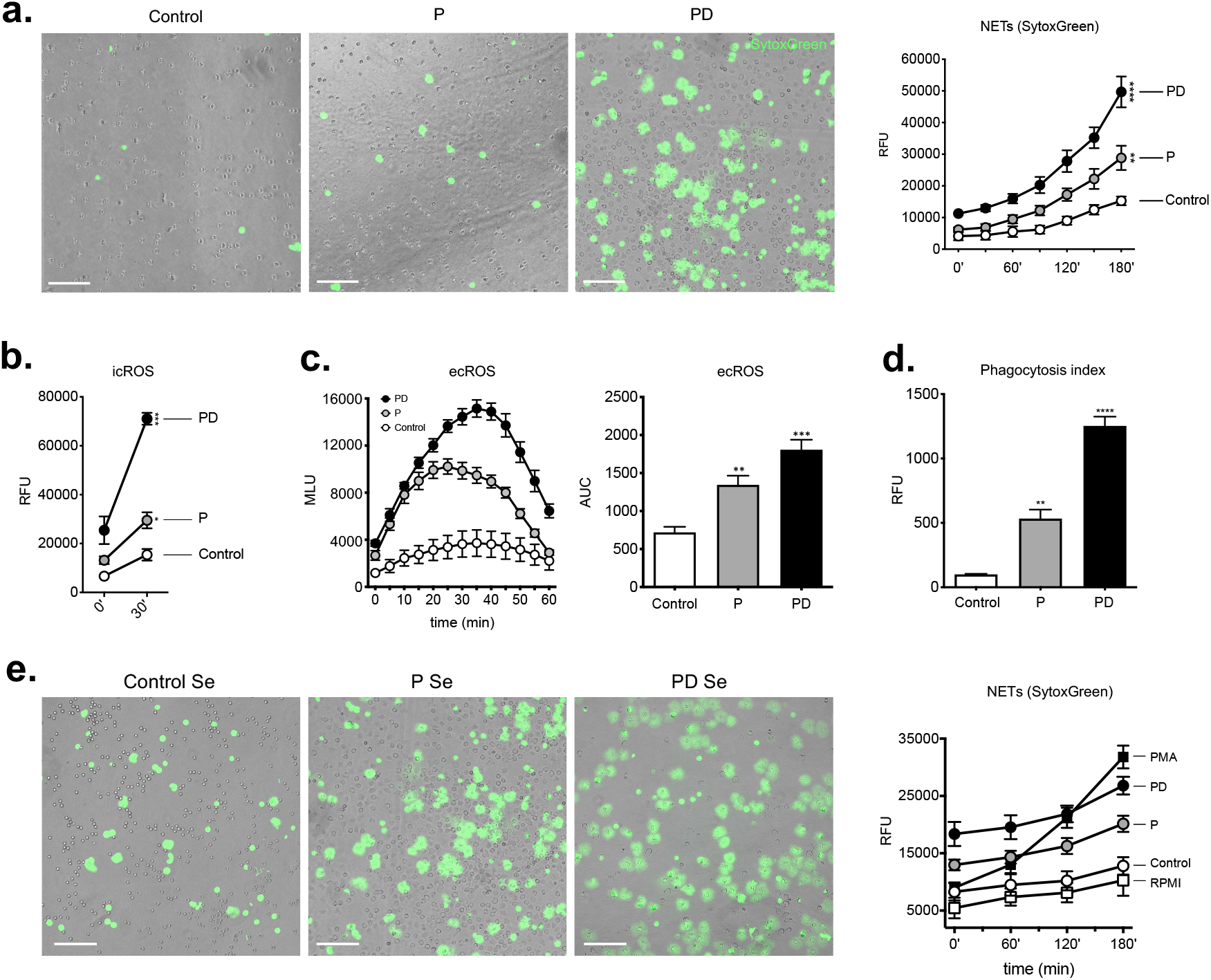
Neutrophil pro-NETotic activity is augmented 48 hours post-delivery. (**a**.) Detection of *in vitro* spontaneous NET formation of neutrophils from healthy controls, donors during pregnancy and postpartum release in a 3-hour time course by fluorescence microscopy using dsDNA-binding fluorescent SytoxGreen dye and fluorimetry (right panel). (**b**.) Intracellular oxidative burst in neutrophils from healthy control donors, donors during pregnancy and postpartum by DCFH-DA detection. (**c**.) Extracellular oxidative burst in neutrophils from healthy control donors, donors during pregnancy and postpartum by luminol assay. (**d**.) Phagocytic activity of neutrophils obtained from healthy female controls, donors during pregnancy and 48 hours post-delivery. (**e**.) *In vitro* spontaneous extracellular DNA release monitored in a 3-hour time course by SytoxGreen dye microscopy after treatment of control neutrophils with pregnancy and post-delivery serum and fluorimetric quantification (right panel). Magnification: 10x. Scale bars: 100 μm. P: pregnancy; PD: post-delivery; ecROS: extracellular ROS; icROS: intracellular ROS; RFU: relative fluorescence units; MLU: mean fluorescence units; AUC: area under the curve. Data are presented as mean ± SEM. ^*^P < 0.05, ^**^P < 0.01, ^***^P < 0.001, ^****^P < 0.0001 (one- or two-way ANOVA followed by Bonferroni’s multiple comparison post-test). All experiments were performed at least 3 times with consistent results.

Simultaneous co-incubation of control naive neutrophils from healthy donors with postpartum serum revealed excessive NET release (Figure 2e) compared to control serum and serum from pregnant women as well, which strongly underlines the augmented pro-NETotic activity of the post-delivery circulatory environment.

### Postpartum circulatory neutrophils are prone to spontaneous *in vitro* NET formation and hyperresponsive to secondary pro-NETotic stimuli

In order to substantiate these observations and further study the cellular responses *per se*, we examined the kinetics of spontaneous NET extrusion in more detail. Neutrophils were cultured for 3 hours and NETs were detected by immunohistochemistry for myeloperoxidase (MPO), a hallmark of NETs imaging and DAPI (4’,6-diamidino-2-phenylindole) counterstaining (Figure 3a). We quantified the degree of *in vitro* NETosis by morphometric analysis and the percentage of area covered by NETs and by citH3-positive intact neutrophils corresponding to neutrophil sensitization towards NET formation (Figure 3b). In parallel, we assessed cell-free nucleosomes in the respective supernatants, specifically their association with myeloperoxidase (MPO), indicative of the NETotic origin of this material (13, 34) (Figure 3c). These experiments indicated that circulating neutrophils derived postpartum generated NETs more rapidly and to a greater magnitude than control and pregnancy neutrophils, a feature evident both at baseline conditions (0h) and at the 3-hour stage (3h) of *in vitro* culture.

**Figure 3.**
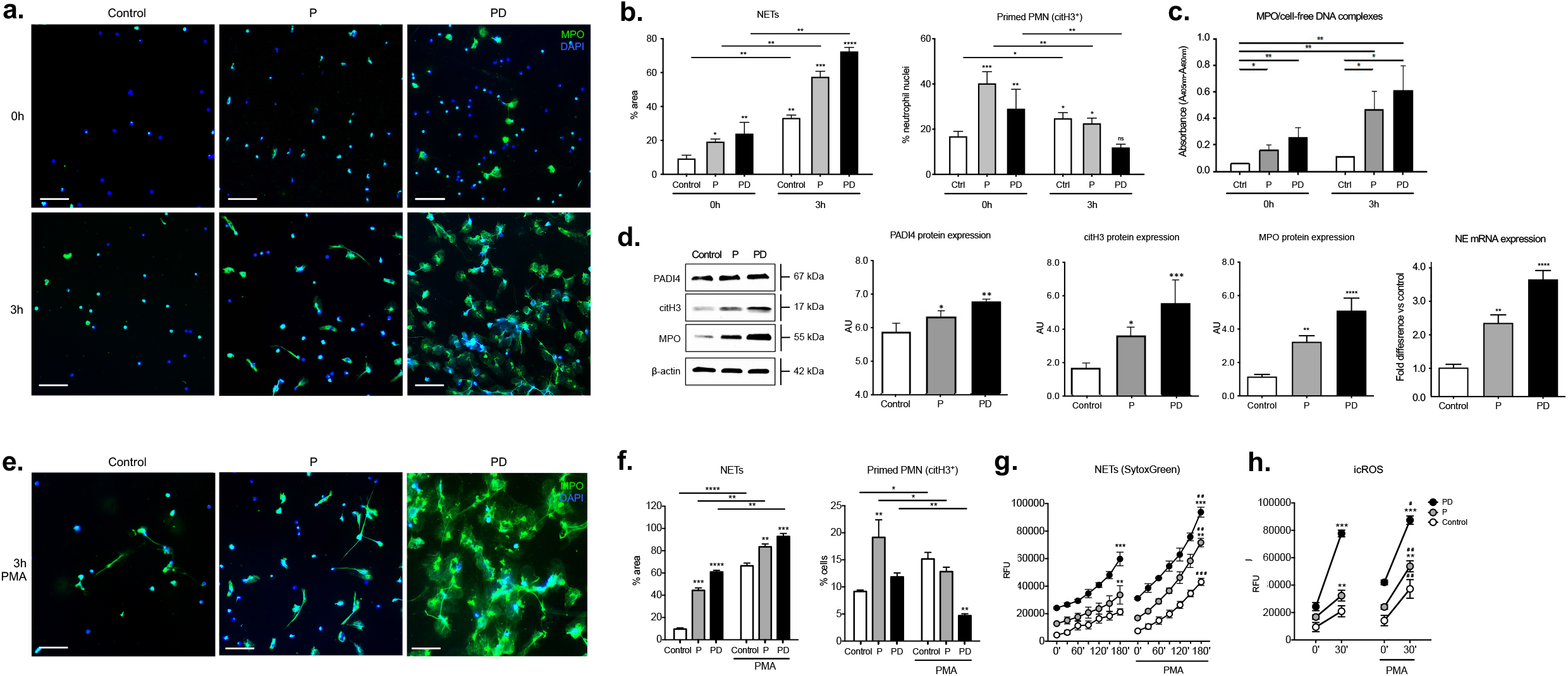
Postpartum circulatory neutrophils are prone to spontaneous *in vitro* NET formation and are hyperresponsive to pro-NETotic stimuli. (**a**.) Fluorescent immunohistochemistry for MPO (green), and DNA counterstain with DAPI (blue) at the baseline initial condition (0h) and after 3 hours in culture (3h). (**b**.) Morphometric analysis of the NETotic (MPO+) and pro-NETotic primed (citH3+) neutrophils from healthy donors, donors during pregnancy and postpartum. (**c**.) Quantification of NETs-associated MPO/DNA complex levels in culture supernatants of neutrophils from healthy controls, donors during pregnancy and postpartum, at the steady state condition (0h) and at the 3-hour timepoint (3h). (**d**.) Western blot and densitometric analysis of PADI4, citH3, MPO and beta-actin protein expression levels in neutrophil lysates from healthy female controls, donors during pregnancy and 48 hours postpartum; NE gene expression analysis by Taqman qRT-PCR in RNA samples obtained from healthy female controls, donors during pregnancy and 48 hours postpartum. (**e**.) Fluorescent immunohistochemistry for MPO (green), and DNA counterstain with DAPI (blue) at the baseline initial condition (0h), after a 3-hour culture (3h). after incubation with the NET-inducing agent horbol-12-myristate-13-acetate (PMA) for a period of 3 hours. Magnification: 20x; Scale bars: 50 μm. (**f**.) Morphometric analysis of the NETotic (MPO+) and pro-NETotic primed (citH3+) neutrophils from healthy donors, donors during pregnancy and postpartum. (**g**.) *in vitro* spontaneous NET formation of neutrophils from healthy controls, donors during pregnancy and postpartum release in a 3-hour timecourse by fluorimetric assessment. (**h**.) Intracellular oxidative burst in neutrophils from healthy control donors, donors during pregnancy and postpartum by DCFH-DA detection. P: pregnancy; PD: post-delivery. Data are presented as mean ± SEM. ^*^P < 0.05, ^**^P < 0.01, ^***^P < 0.001, ^****^P < 0.0001 (one-way ANOVA followed by Bonferroni’s multiple comparison post-test). All in vitro experiments were performed at least 6 times with consistent results.

NETosis has been shown to depend on several biochemical signalling elements, among which are the generation of reactive oxygen species (ROS) by nicotinamide adenine dinucleotide phosphate (NADPH) oxidase, the actions of MPO and NE, as well as histone citrullination by PAD4 (10, 34, 35). In this context, postpartum circulating neutrophils exhibited a marginal increase in PAD4 levels in total cellular protein extracts compared to the control neutrophils, as shown by western blot analysis (Figure 3d). This was associated with a more pronounced increase in citrullinated histone H3 (citH3), MPO and NE mRNA levels when compared to controls (Figure 3d).

Finally, it was noted in our experiments that when postpartum-derived neutrophils were treated with phorbol ester (PMA), they formed NETs far more vigorously than control neutrophils (Figure 3e), which was confirmed by morphometric analysis (Figure 3f), SytoxGreen kinetics quantification (Figure 3g) and ROS release response assessment (Figure 3h). These findings demonstrate a well-defined neutrophil pro-NETotic phenotype postpartum.

### Postpartum NET formation is independent of G-CSF

Circulating G-CSF levels gradually increase during gestation (36), and render neutrophils pro-NETotic (21, 37, 38). Notably, serum G-CSF was highly elevated in postpartum samples, a trend evident to a lesser extent in the respective plasma samples (Figure 4a). To evaluate whether plasma G-CSF could be responsible for the augmented tendency of peripheral blood neutrophils to form NETs, we pre-treated pregnancy and post-delivery plasma with a G-CSF neutralizing antibody and applied it on control neutrophils *in vitro*. In this experiment, pre-treatment with pregnancy plasma containing neutralizing antibody prevented neutrophil sensitization towards NET generation (Figure 4b and Figure 4c, upper panels) and reduced their capacity to generate ROS (Figure 4d, upper panels). Immunostaining showed hypercitrullination of H3, consistent with a predisposition of neutrophils exposed to G-CSF to form NETs (Figure 4e, upper panel and Figure 4f).

**Figure 4.**
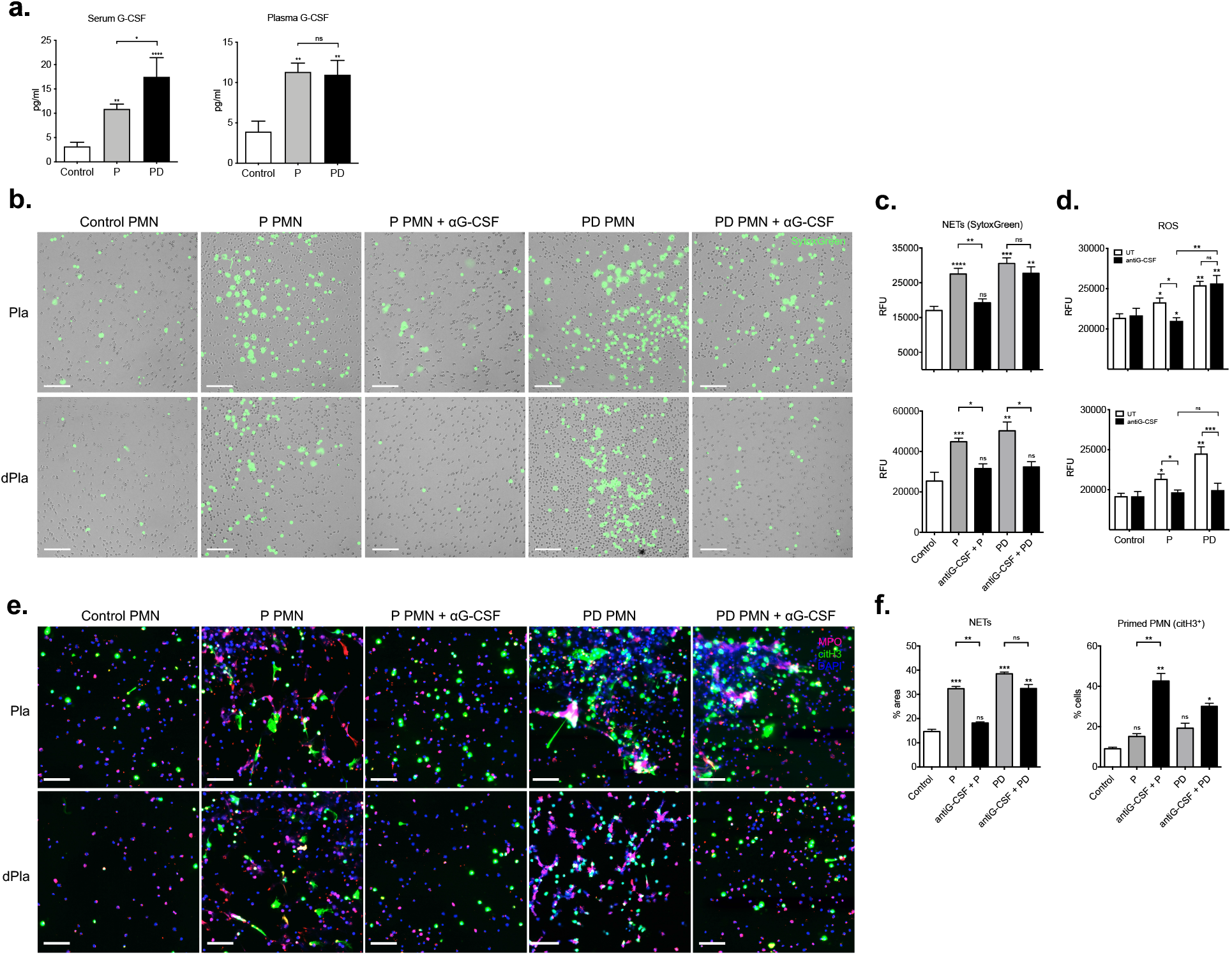
Postpartum NET formation is independent of G-CSF. (**b**.) serum and plasma concentrations of G-CSF in healthy non-pregnant controls, pregnant individuals and at post-delivery. (**b**.) NET formation assessed microscopically in a 3-hour time course with SytoxGreen DNA binding dye after *in vitro* incubation of control neutrophils with pregnancy and postpartum plasma (upper panel) and microparticle-depleted plasma with and without antiG-CSF neutralizing antibody pretreatment. Magnification: 10x. Scale bars: 100 μm. (**c**.) Fluorimetric evaluation of extracellular DNA release and (**d**.) measurement of ROS generation by the DCFH-DA plate assay in control neutrophils treated with pregnancy and post-delivery plasma (upper panels) or depleted plasma (lower panels) in the presence of antiG-CSF neutralizing antibody. (**e**.) Fluorescent immunostainings for MPO (red), citH3 (green) and DNA (blue) and (**f**.) Morphometric analysis of the NETotic (MPO+) and pro-NETotic primed (citH3+) neutrophils from healthy donors, donors during pregnancy and postpartum. Magnification: 20x; Scale bars: 50 μm. P: pregnancy; PD: post-delivery; aG-CSF: antiG-CSF neutralizing antibody. Data are presented as mean ± SEM. ^*^P < 0.05, ^**^P < 0.01, ^***^P < 0.001, ^****^P < 0.0001 (one way ANOVA followed by Bonferroni’s multiple comparison post-test). All experiments were performed at least 6 times with consistent results.

Unexpectedly, G-CSF neutralization was ineffective in post-delivery plasma. Nevertheless, NET formation was still increased, as was the pro-NETotic phenotype, suggesting the presence of an additional factor acting as a secondary stimulus for NETosis in the postpartum plasma. In consideration that NET formation might be driven by placental microparticles (19, 39, 40), we repeated the aforementioned experimental procedure with microdebris-depleted plasma (dPla). Pre-treatment of postpartum plasma with both the G-CSF neutralizing antibody pregnancy and microparticle depletion abolished NET generation (Figure 4b and Figure 4c, lower panels) and reduced ROS generation (Figure 4d, lower panels). Moreover, immunostaining revealed enhanced H3 citrullination in both cases, confirming the postpartum neutrophil pro-NETotic phenotype (Figure 4e and Figure 4f, lower panel).

### Excessive NET formation by postpartum circulating neutrophils is complemented by coagulation cascade activation, platelet activation status alterations and the involvement of circulating microparticles

Since pregnancy represents a *de facto* pro-coagulant state (41, 42), and since NETs have been implicated in pro-thrombotic coagulopathies (17, 37), we sought to examine such features in our study cohort. Indeed, a seminal observation in the context of clotting activation is that TF, a central triggering element of the extrinsic pathway of coagulation, was detected by immunohistochemistry decorating the NETs following *in vitro* culture (Figure 5a), which was also highly pronounced on NETs from post-delivery samples. Moreover, in these examinations we observed that the total number of platelets measured in the whole EDTA blood samples tends to decline during pregnancy, reaching the lowest levels 48 hours postpartum compared to the controls (Figure 5b). This depletion of platelets during pregnancy and especially postpartum is associated with a change in their physical properties, as determined by parameters such as mean platelet volume (MPV) and platelet distribution width (PDW). These features potentially reflect upon the platelet activation status, with platelets having a high MPV tending to promote coagulation (43) (Figure 5c), and may be of particular significance, as neutrophil NETs can be intensively induced by activated platelets (44). Since the MPV increases after delivery, this could indicate functional, metabolical, and enzymatical activity and an escalating chemokine and cytokine production (45). Furthermore, this change in platelet volume is mirrored to a lower extent by the PDW, which was again significantly elevated 48 hours postpartum (Figure 5c), indicating clotting activity associated with delivery (46).

**Figure 5.**
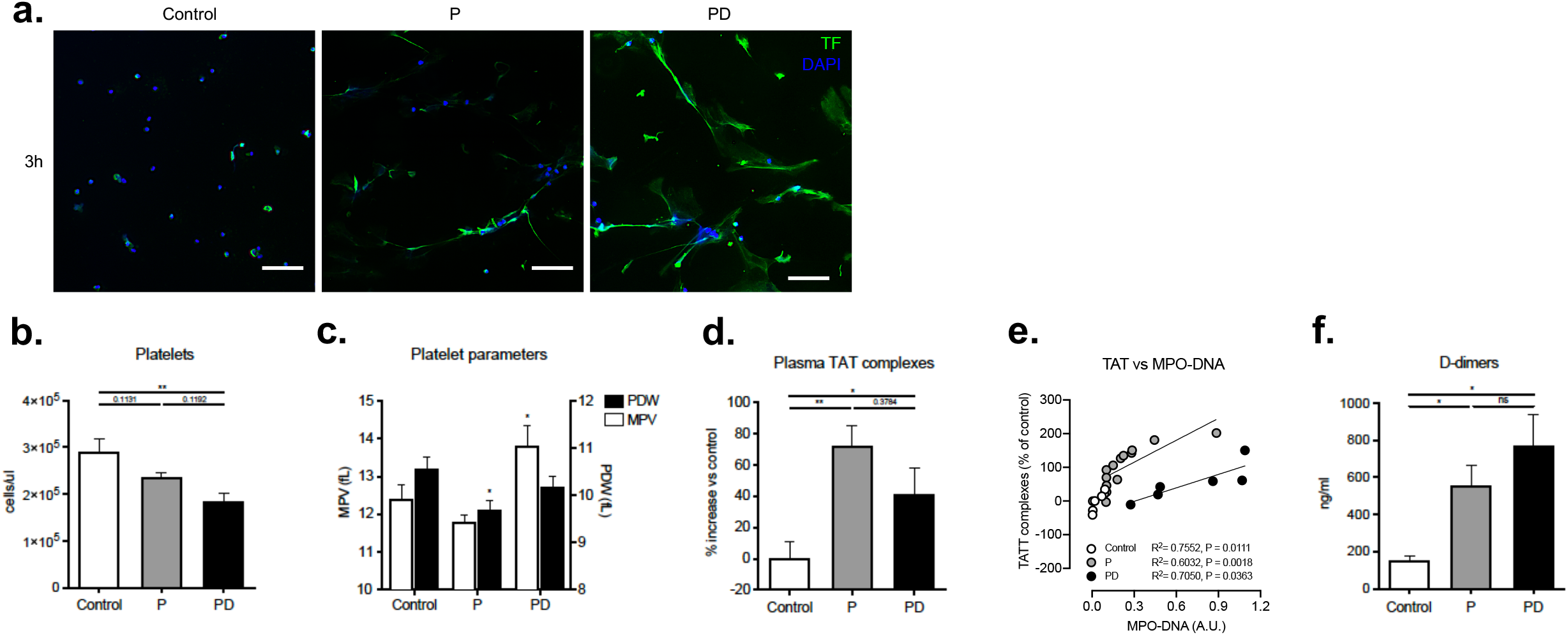
Excessive NET formation by postpartum circulating neutrophils is complemented by activation of the coagulation cascade and alterations in the platelet activity status. (**a**.) Immunohistochemical staining for TF (green), and DNA counterstain with DAPI (blue) after a 3-hour *in vitro* culture. Magnification: 20x; Scale bars: 50μm. (**b**.) Platelet differential counts in healthy non-pregnant blood donors, donors during pregnancy and 48 hours postpartum. (**c**.) Comparative analysis between healthy donor controls, donors during gestation and postpartum, concerning the platelet morphology and activation indices mean platelet volume (MPV) and platelet distribution width (PDW). (**d**.) TAT complexes concentration in citrate plasma from healthy donor controls, donors during pregnancy and postpartum, determined by ELISA. (**e**.) Spearman correlation between TAT complexes concentration and NET-associated MPO/DNA complexes. (**f**.) D-dimer levels in citrate plasma from healthy donor controls, donors during gestation and postpartum, determined by ELISA. P: pregnancy; PD: post-delivery. Data are presented as mean ± SEM. ^*^P < 0.05, ^**^P < 0.01, ^***^P < 0.001, ^****^P < 0.0001 (one way ANOVA followed by Bonferroni’s multiple comparison post-test). All experiments were performed at least 6 times with consistent results.

We also measured other clotting parameters such as thrombin-antithrombin (TAT) complexes and D-dimer levels. The high levels of free TAT complexes and D-dimers detected during pregnancy suggest that gestation is associated with an active clotting cascade leading to thrombin formation and fibrinolysis (Figure 5d and Figure 5f). Of note, however, is that there seems to be a difference in the kinetics between these two factors in postpartum samples, with TAT complex levels decreasing compared to gestation but still higher than in the healthy controls, while D-dimer concentrations further increase in early postpartum (Figure 5d and Figure 5f). This is also reflected in the significant correlation between TAT complexes and NETs where pregnancy and postpartum curves exert similar but distinctive tendencies (Figure 5e). These differences may reflect the two different coagulation parameters assessed by these two assays, in that TAT complexes are indicative of de-activated thrombin, whereas D-dimers are suggestive of prior thrombin activity or blood clot fibrinolysis.

Finally, microparticles obtained from the plasma depletion process were quantified and used for *in vitro* stimulation studies (Fig. 6a). After stimulation, considerably higher levels of cell-free DNA and NETs, assessed by SytoxGreen fluorimetry and imaging, were recorded, with the peak levels reached by the postpartum plasma MPs (Fig. 6b). ROS generation measurements complemented these observations (Fig. 6c), as well as the immunohistochemical staining and morphometric analysis (Fig. 6d and Fig. 6e). Interestingly, the citH3 positive pro-NETotic phenotype was still evident only in neutrophils treated with pregnancy plasma MPs, but not with the postpartum plasma micro-debris (Fig. 6d). Collectively, these data suggest a well-defined qualitative, beyond the quantitative, distinction between pregnancy and post-delivery plasma-derived microparticles, which exacerbate the NET forming capacity in the complete absence of sex hormones after placental expulsion at term.

**Figure 6.**
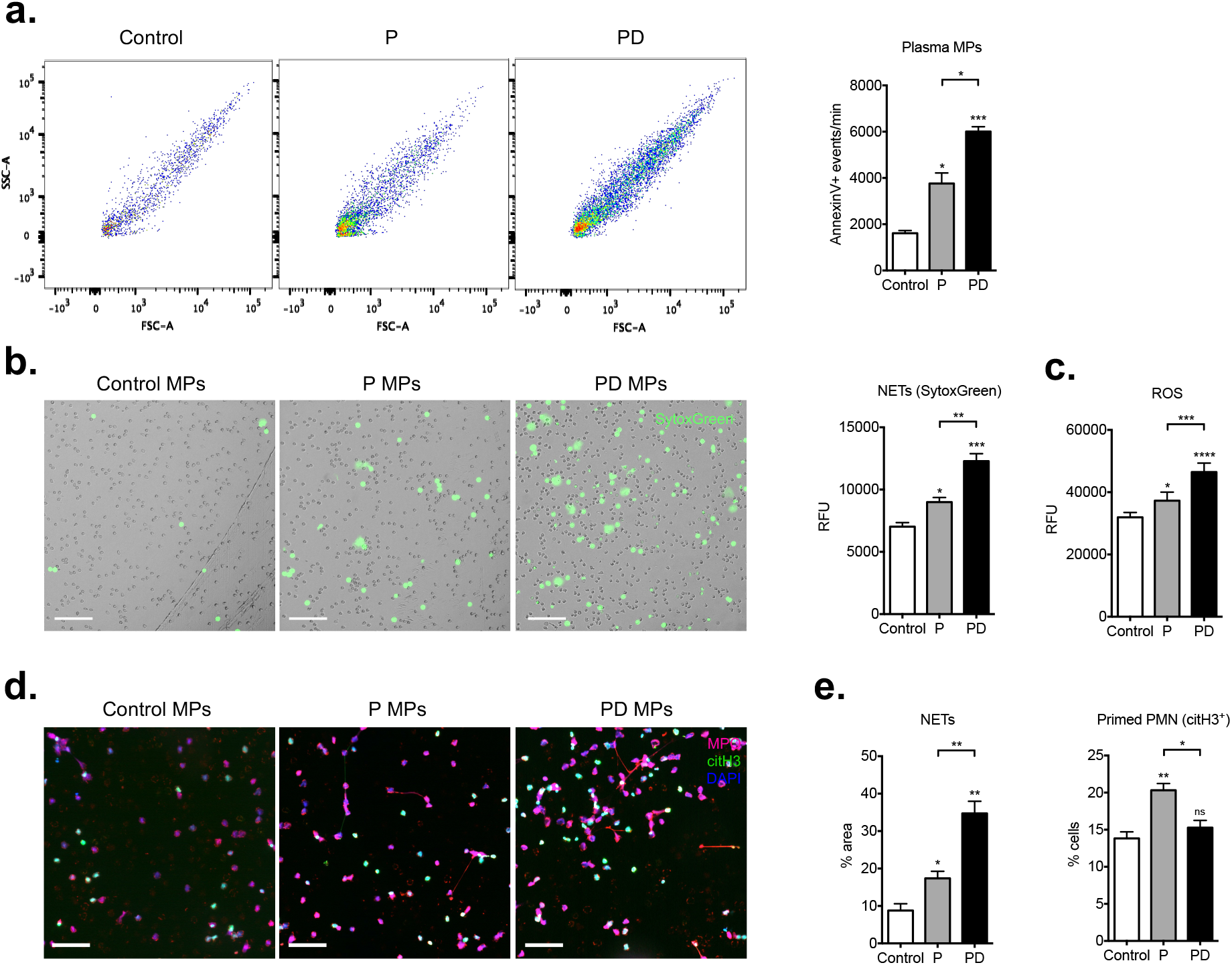
Postpartum NET formation is triggered by circulating microparticles. (**a**.) Fluorescence histograms and bar graph depicting the quantitative analysis of microparticles in control, pregnancy and 48 hours postpartum plasma, positive for AnnexinV staining. Plasma was assessed for 60 seconds. (**b**.) *In vitro* NET formation detected microscopically in a 3-hour time course with SytoxGreen DNA binding dye after a 1-hour pretreatment with equal amounts of microparticles isolated from pregnancy plasma and 48 hours postpartum plasma and SytoxGreen fluorimetry of *in vitro* extracellular DNA release by control neutrophils (right panel) after 3 hours treatment with pregnancy and post-delivery microparticles. (**c**.) Measurement of ROS generation by control neutrophils in response to pregnancy and postpartum derived microparticles by DCFH-DA detection. (**d**.) Fluorescent immunostainings for MPO (red), citH3 (green) and DNA (blue) and (**e**.) morphometric assessment of NETs (MPO positive) and pro-NETotic (citH3 positive) neutrophils under the same experimental setup. Magnification: 20x; Scale bars: 50 μm. P: pregnancy; PD: post-delivery. Data are presented as mean ± SEM. ^*^P < 0.05, ^**^P < 0.01, ^***^P < 0.001, ^****^P < 0.0001 (one-way ANOVA followed by Bonferroni’s multiple comparison post-test). All experiments were performed at least 3 times with consistent results.

## Discussion

The post-delivery period represents a highly critical phase for the survival of the mother (1). A completely new immunological environment is formed rapidly after childbirth, in which the immune system and neutrophils in particular need to respond efficiently to a highly perilous situation by mounting sufficient haemostasis, enhanced protection against infection and effective wound healing. It is not surprising that postpartum haemorrhage, puerperal infections and venous thromboembolism (VTE) are still associated with high maternal mortality and morbidity (1, 5, 6). The role of neutrophils is essential in these processes, given that disturbed neutrophil adaptation represents a detrimental cause for the abovementioned complications.

We hereby demonstrate that, shortly after delivery, a systemic milieu is shaped that sensitizes peripheral blood neutrophils towards a pro-NETotic state, rendering them able to readily form NETs upon a secondary effector signal provided by secretory microparticles. This mechanism seems applicable to the general immunological context of the puerperium. Throughout pregnancy, as the levels of sex hormones and inflammatory cytokines rise, the maternal immune system balances opposing requirements. On the one hand, tolerance of the semi-allogeneic fetus sustains its integrity, while on the other maintenance of a robust immune reactivity protects both mother and fetus from invading pathogens with neutrophils playing a pivotal role (2-4). During pregnancy neutrophils are primed to mount an efficient anti-microbial defense by deploying NETs, contributing critically to feto-maternal well-being (39). Sex hormones, including hCG, estrogens and progesterone, contribute largely to successful pregnancy by modulating the immune response. Numerous studies to date support the impact of the endocrine adaptations in the regulation of chemotactic neutrophil recruitment and neutrophil-endothelial cell interaction (2, 3, 47, 48). Since postpartum neutrophils displayed an enhanced NET formation when treated with PMA, a basal primed pro-NETotic state, also observed during pregnancy, may be assumed (20, 21, 39).

Consistent with our findings, the propensity to form NETs peaks early post-delivery, when the risk of maternal infection and morbidity/mortality is greatest and the need for an additional clot propagating and stabilizing mechanism is imminent. Alongside the enhanced phagocytic activity and degranulation, our morphometric analysis indicated that phenotypically quiescent neutrophils during puerperium possess a distinctive nuclear morphology, characterized by a delobulated nuclear and citH3 positive phenotype. This special pro-NETotic sensitization of the postpartum neutrophils develops in an environment where sex hormones, which finely regulates the balance towards NET formation during pregnancy, are totally absent due to the expulsion of the placenta (39, 49-52).

G-CSF plays a major role in maintaining basal neutrophil homeostasis through the regulation of neutrophil release from the bone marrow to the circulation (7, 8). It’s importance, however, in modulating neutrophil activity is less clear. Several reports point to a certain role in regulating specific aspects of the overall immune response, innate and adaptive, to bacterial clearance and local inflammation during infections (53, 54). Interestingly, the mobilization of neutrophils is partially driven by NE (55). Since elevations of circulatory G-CSF lead to increased numbers of neutrophils (7, 53), but also promote pro-NETotic priming in animal model systems (37), we postulated that such a system is operative not only in pregnancy but also postpartum. Our data confirm and extend upon previous reports regarding elevated basal serum levels of G-CSF (39). As gestation progresses, pro-NETotic activity becomes modulated by locally secreted G-CSF and shed microparticles from cells and tissues to proximate neutrophils. In this regard, elevated concentrations of G-CSF early postpartum not only serve to increase the pool of circulatory neutrophils, but also to promote a primed pro-NETotic phenotype. This is accomplished by triggering increased PAD4, MPO and NE expression and concomitant histone H3 citrullination, essential steps in the signalling cascade leading to formation of NETs (35, 56). Moreover, to meet the enormous demand for neutrophils during infection- or malignancy-related inflammation, G-CSF-driven steady-state granulopoiesis is switched to emergency granulopoiesis, which is characterized by considerably enhanced *de novo* generation of neutrophils, accelerated cellular turnover and the release of immature and mature neutrophils from the bone marrow into the peripheral blood (57, 58). Excessive G-CSF as the key signal can be locally generated by non-hematopoietic cells and tissues too, such as the activated vascular endothelium (57), with increased levels of vWF, elevated thrombin generation and an eminent procoagulant state (22). This correlates with previous observations claiming that postpartum granulocytes possess characteristics of less mature neutrophils compared to cells from healthy control donors, and appear to be more prone to secondary stimuli (59, 60). The phenotypic and functional features of postpartum neutrophils identified and characterized in the present study support the concept of emergency granulopoiesis.

Pregnant women are confronted with a fivefold higher risk for venous thrombosis (6), which is dramatically exacerbated in the occurrence of pre-eclampsia (PE) (61). In similar settings, NETs generated by primed neutrophils could lead to enhanced thrombosis and inflammation in autoimmune or virally-induced vasculitis by promoting the expression of TF, the initiator of the extrinsic pathway of the coagulation cascade (18, 62, 63), while the involvement of activated platelets in thrombo-inflammation is indispensable (64, 65). Conversely, thrombi *per se* contain neutrophils and NETs that are decorated with TF (16), a process shown to be largely governed by IL-1β (66). Intriguingly, the threat of thrombotic embolism peaks immediately after delivery (6, 62). In normal pregnancy, increased venous thrombotic embolism seems to progress due to a *de facto* gestational hypercoagulative state, a mechanism utilized to protect from blood loss during childbirth (6, 67, 68). The observation that NETs promote coagulation, thereby contributing to clot formation, sheds new light into this process (17, 35, 37). The exact means whereby NETs promote coagulation is currently under investigation, but likely includes interaction with activated platelets (69-71). If left unchecked, this could initiate a feedback loop with potentially disastrous consequences.

Furthermore, it seems that intact NETs may not be essential for active clot formation, but rather that individual NET components such as cell-free DNA, histones or liberated serine proteases provide the crucial momentum (70, 72). Previous evidence indicates that PE is associated with increased concentrations of circulatory cfDNA (21), which provides vital clues for the increased incidence of venous thrombotic embolism during pregnancy (33). Additional support for such a supposition comes from the fact that the concentrations of maternal cfDNA are highest during labour and immediately postpartum, and that they were significantly elevated in cases with severe PE (73). Should these factors indeed contribute to enhanced coagulation, our data fit well with the increased incidence of venous thrombotic embolism under such conditions, but possibly also shortly after parturition (33).

The prevalence and the progression of several autoimmune disorders are affected by hormonal fluctuations (4, 47, 74). In contrast, a gestation-related decline in disease activity is experienced in a majority of autoimmune disease patients,with flares recurring after delivery (75, 76). Neutrophils and NETosis appear to play an important role in the pathogenesis of the above mentioned disorders (12, 76, 77). With regard to pregnancy, it is plausible that in extreme conditions such as PE, NETosis additionally promotes a strong pro-coagulant state, possibly resulting in placental infarction (40, 78). Strikingly, maternal sex hormone levels in the circulation drop dramatically immediately after removal of the placenta at term (49-52). Moreover, a vast influx of cell- and tissue-derived microparticles is observed in the postpartum circulation, conveying pro-inflammatory, immune activating, and pro-coagulant activities (73, 79). Our analyses suggest that pregnancy serum and plasma, which contain vast amounts of diverse microdebris of mixed background, lead to excessive NET formation depending on the microparticle load. The possibility for such a functional connection is also supported by our own observation that while depleted pregnancy plasma fails to induce NETs, the respective isolated microparticles effectively promote excessive NETosis *in vitro*, which strongly correlates with previous findings regarding PE (19, 79, 80). Moreover, previous studies showed that tumour-derived microparticles induce NETs in neutrophils from mice treated with G-CSF and interacted with NETs under static conditions (81). Accordingly, the intravenous administration of tumour microparticles into G-CSF-treated mice significantly accelerated venous thrombosis *in vivo*, suggesting that microparticles and neutrophils cooperate in establishing cancer-associated thrombosis (81).

In the same line of thought, microorganisms such as viruses are known to activate platelets to shed microparticles (82), which in turn further activate neutrophils and macrophages, thereby inducing formation of NETs and proinflammatory cytokine release in a detrimental feedback loop (82). Hence, the enhanced NETotic capacity noted in post-delivery neutrophils, which could not be neutralized by the anti-G-CSF antibody, contributes to a rather robust pro-coagulant milieu. This type of involvement might act primarily in a protective fashion, diminishing the risk of severe haemostatic and infection-related postpartum complications. These aspects need to be addressed, though, in detail in forthcoming studies.

In summary, the presented data demonstrate that early after delivery neutrophils exhibit a pro-NETotic state and an enhanced propensity to release NETs in response to specific physiologic stimuli. This activity is modulated at several key levels. First, G-CSF is a major signal, inducing neutrophil release from the bone marrow, providing an important mechanism by which the level of circulating neutrophils and their sensitization is increased. Second, the degree of pre-activation seems to be finely tuned by the quantity of hormones and microparticles, which are produced by a variety of tissues, including the injured endothelium, and reach their peak concentrations shortly after term. Finally, TF-enriched NETs are released locally in proximity to damaged endothelium, where additional players - such as activated platelets and polymerized chains of von Willebrand Factor (vWF) - might already deploy under the presence of excessive post-delivery secondary stimuli (Fig. 7). This regulatory mechanism of coagulation activation and thrombus stabilization may underlie changes in susceptibility to postpartum complications due to harmful NET components. Our findings regarding neutrophil responses in puerperium provide new insight concerning prothrombotic gestation-related pathologies, since neutrophil recruitment, activation and NET release could be associated with excessive endothelial and placental injury, leading the path for effective complementary pharmaceutical interventions.

**Figure 7.**
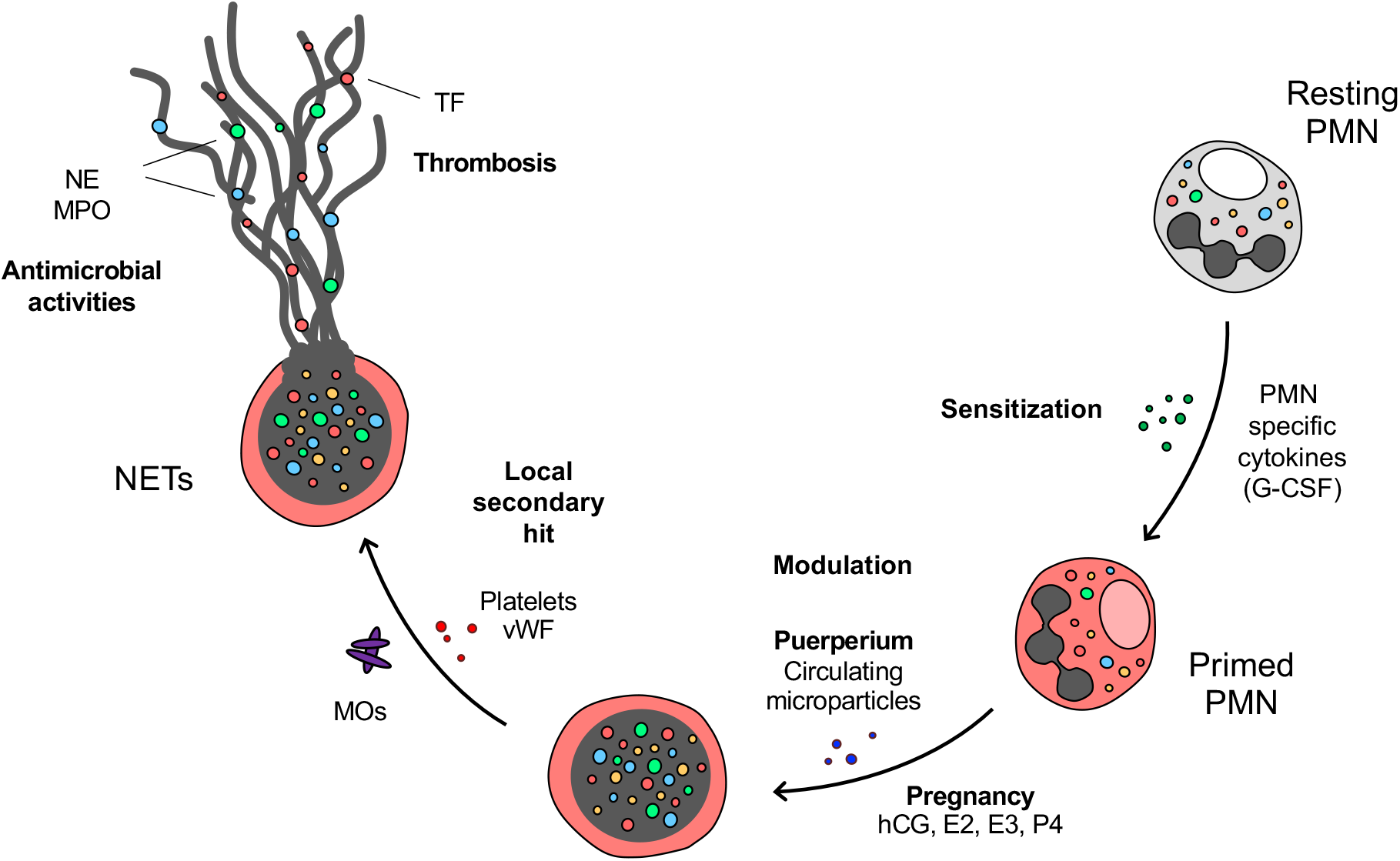
Schematic depiction of the mechanism by which neutrophils exhibit a pro-NETotic state and enhanced tendency to release NETs early after delivery in response to specific physiologic stimuli promoting thrombosis. G-CSF is a major signal inducing neutrophil release from the bone marrow, providing an important emergency mechanism by which the level of circulating neutrophils and sensitization is increased postpartum. The degree of neutrophil activation seems to be finely tuned by the unique stoichiometry of surrounding sex hormones and microdebris, which are produced by a variety of tissues and cells, including the wounded endothelium and platelets, and reach their peak concentrations shortly after term. Finally, NETs are released locally after this second hit in the vicinity of activated endothelial loci, where polymerized chains of von Willebrand Factor (vWF) might already deploy under the presence of excessive post-delivery secondary stimuli, thus facilitating the thrombotic cascade.

## Abbreviations

(PMN): Polymorphonuclear granulocytes
(NETs): neutrophil extracellular traps
(DVT): deep vein thrombosis
(RA): rheumatoid arthritis
(citH3): citrullinated H3
(NE): neutrophil elastase
(MPO): myeloperoxidase
(TAT): thrombin-antithrombin complexes
(G-CSF): granulocyte colony stimulating factor
(DAPI): 4’,6-diamidino-2-phenylindole
(PMA): phorbol ester
(ROS): reactive oxygen species
(NADPH): nicotinamide adenine dinucleotide phosphate oxidase
(citH3): citrullinated histone H3
(MPV): mean platelet volume
(PDW): platelet distribution width
(TF): tissue factor
(SLE): systemic lupus erythematosus
(PE): preeclampsia
(PAD4): peptidylarginine deiminase 4
(MP): microparticles

## Author Contributions

SG, CC, MS and GS performed all experiments; SVB, AB, IH and OL provided advice for and contributed to the recruitment of the sample donors; PH, UAW and SH provided advice and contributed to the revision of the manuscript; SG and SH devised and directed the study; and SG wrote the manuscript.

## Acknowledgments

SG was supported by grants from the Fonds W + W and the Research Council of Kantonsspital Aarau to PH. MS was supported by a grant from the Swiss Red Cross Basel to SH. We thank Alina Wunderle for initial observations, Umabalini Nagalingam for her assistance with western blot analysis, and Giuliano Bayer for his assistance with neutrophil imaging. Finally, we wish to thank Andre Tiaden, Basel, BS, CH; Marko Radic, Memphis, TN, USA; and Venizelos Papayannopoulos, London, UK, for helpful discussions and their comments on the manuscript.

