## Supplemental Table 1 for "Circulatory neutrophils exhibit enhanced neutrophil extracellular trap formation in early puerperium: NETs at the nexus of thrombosis and immunity?"

| | Non-pregnant | Pregnant | | Postpartum | P |
|---|---|---|---|---|---|
| n | 20 | | 29 | 9 | na |
| | | IT: | 5 | | |
| | | IIT: | 13 | | |
| | | IIIT: | 11 | | |
| Maternal age (years) | 32.7 (23-46) | 33.9 (26-41) | | 38.2 (25-40) | ns |
| Gestational age (weeks) | na | IT: | 12.5 (12-13) | na | na |
| | | IIT: | 24.7 (22-26) | | |
| | | IIIT: | 39.1 (37-41) | | |

Values expressed as mean ± SEM with minimum-maximum, where applicable; IT, first trimester; IIT, second trimester, IIIT, third trimester ns, not significant; na, not applicable.

**Table S1.** Demographic characteristics of the study cohort, i.e. women during the three trimesters of pregnancy and non-pregnant healthy controls.
